# Structure and function of the human mitochondrial MRS2 channel

**DOI:** 10.1101/2023.08.12.553106

**Authors:** Zhihui He, Yung-Chi Tu, Chen-Wei Tsai, Jonathan Mount, Jingying Zhang, Ming-Feng Tsai, Peng Yuan

## Abstract

The human Mitochondrial RNA Splicing 2 protein (MRS2) has been implicated in Mg^2+^ transport across mitochondrial inner membranes, thus playing an important role in Mg^2+^ homeostasis critical for mitochondrial integrity and function. However, the molecular mechanisms underlying its fundamental channel properties such as ion selectivity and regulation remain unclear. Here, we present structural and functional investigation of MRS2. Cryo-electron microscopy structures in various ionic conditions reveal a pentameric channel architecture and the molecular basis of ion permeation and potential regulation mechanisms. Electrophysiological analyses demonstrate that MRS2 is a Ca^2+^-regulated, non-selective channel permeable to Mg^2+^, Ca^2+^, Na^+^ and K^+^, which contrasts with its prokaryotic ortholog, CorA, operating as a Mg^2+^-gated Mg^2+^ channel. Moreover, a conserved arginine ring within the pore of MRS2 functions to restrict cation movements, likely preventing the channel from collapsing the proton motive force that drives mitochondrial ATP synthesis. Together, our results provide a molecular framework for further understanding MRS2 in mitochondrial function and disease.

## Introduction

Mg^2+^, the most abundant divalent cation in living organisms, plays critical physiological roles by forming the biologically active Mg^2+^-ATP complex, modulating chromatin stability, and regulating enzyme activity. Abnormal cellular Mg^2+^ levels have been linked to a wide range of disorders in the cardiovascular, muscular, nervous, digestive and immune systems^1^. In mammalian cells, Mg^2+^ homeostasis is managed by an array of Mg^2+^ channels and transporters, including TRPM6/7, SLC41A1, CNNM, MagT1 and MRS2 (mitochondrial RNA splicing 2)^2^. Accumulating studies have indicated that MRS2, which belongs to the superfamily of CorA Mg^2+^ transporters that are found in organisms including bacteria, fungi, plants and animals^3^, is the molecular conduit that imports cytosolic Mg^2+^ into the mitochondrial matrix to regulate mitochondrial metabolism and inter-organellar communication^4–9^. Dysfunction or downregulation of MRS2 has been associated with severe pathologies, including neurodegeneration, depressed oxidative-phosphorylation, reduced cell viability and aggravated cancer progression^10–16^.

Despite the critical roles of MRS2 in mammalian physiology and pathophysiology, understanding of its molecular mechanisms remains limited and is primarily inferred from studies of the distantly related prokaryotic CorA Mg^2+^ channels^2,17^. Structural studies of *Thermotoga maritima* (TmCorA) and *Methanocaldococcus jannaschii* CorA (MjCorA) have revealed a pentameric assembly with a central ion permeation pathway, consistent with an ion-channel architecture^18–21^. Moreover, the signature Gly-Met-Asn (GMN) motif is proposed to function as the ion selectivity filter, with the side chains of five asparagine residues generating a pentameric polar ring for high-affinity Mg^2+^ coordination at the extracellular pore entrance^18–21^. Electrophysiological characterization further demonstrates that TmCorA exhibits the anomalous mole fraction effect (AMFE), diagnostic of the classical single-file, multi-ion-pore conduction mechanism^22,23^. Additional Mg^2+^-binding sites, identified at the cytoplasmic interfaces between neighboring subunits, confer Mg^2+^-dependent gating^23–25^. Specifically, the absence of Mg^2+^ at these interfacial sites favors a conductive conformation. As Mg^2+^ influx via CorA raises the local Mg^2+^ concentration, subsequent binding of Mg^2+^ to these cytoplasmic sites would induce channel inactivation, analogous to how Ca^2+^ influx inactivates voltage-gated Ca^2+^ channels^26,27^.

While it is tempting to infer from these studies that mammalian MRS2 also functions as a Mg^2+^-gated Mg^2+^ channel, the low sequence identity between CorA and MRS2 (<20%) raises a possibility of significant divergence in their channel properties. Indeed, quantitative ^28^Mg flux assays have demonstrated puzzlingly slow kinetics of mitochondrial Mg^2+^ uptake^28,29^, >100-fold slower than a mitochondrial Ca^2+^ channel known as the mitochondrial Ca^2+^ uniporter^30,31^. This suggests that MRS2 might have a much smaller Mg^2+^ conductance than CorA or that MRS2 is subjected to tight regulation in the mitochondrial inner membrane. Moreover, the finding that another prokaryotic CorA family member, ZntB, functions as a proton regulated Zn^2+^ transporter^32^, indicates that proteins with the CorA structural fold can evolve to transport a range of different cations via distinct regulation mechanisms to fulfill particular physiological purposes. These uncertainties prompted us to launch a detailed investigation combining electrophysiology and single-particle cryo-electron microscopy (cryo-EM) to examine the fundamental properties of MRS2, including its molecular structure, ion permeation and channel regulation mechanisms. Determination of these basic channel properties is essential to understand how MRS2 regulates cellular physiology and provides a foundation to modulate MRS2 activity to alleviate pathological conditions.

## Results

### Structure determination

Heterologous expression of the full-length human MRS2 protein (443 amino acids) was insufficient for structural studies. Guided by the AlphaFold prediction^33^, we constructed a truncated channel core consisting of amino acids 62-431, in which the presumably unstructured N- and C-terminal portions were removed. However, this engineered construct showed marginal improvement in overexpression. This motivated us to apply a fusion strategy, in which a thermostabilized BRIL protein, proven valuable to enhance expression and stability of suboptimal membrane proteins^34–37^, was attached to the C-terminal end of MRS2. The resulting construct, dubbed MRS2_EM_, expressed well in yeast *P. pastoris* and showed excellent biochemical properties suitable for structural studies.

To establish the molecular basis of ion recognition and conduction in MRS2, we conducted single-particle cryo-EM experiments under several different ionic conditions. With the inclusion of 10 mM ethylenediaminetetraacetic acid (EDTA) to chelate Mg^2+^ or Ca^2+^ ions in the protein purification buffer, we obtained the MRS2_EDTA_ structure that is presumably free of divalent ions (Extended Data Fig. 1). To investigate Mg^2+^ and Ca^2+^ permeation, we determined the cryo-EM structures in the presence of 40 mM Mg^2+^ (MRS2_Mg_) or 10 mM Ca^2+^ (MRS2_Ca_) (Extended Data Fig. 1). These structures, with resolutions ranging from ∼2.9 to 3.1 Å, are of good quality and enable unambiguous model building (Extended Data Fig. 2, Extended Data Table 1). Comparison of these structures reveals that the MRS2 channel maintains an essentially identical conformation in these various ionic conditions, but ion occupancy differs in these structures.

### Overall architecture

Human MRS2 forms a pentameric channel with a central ion conduction pore along the five-fold symmetry axis (Fig. 1a, b). In each subunit, the N-terminal region, known to dwell within the mitochondrial matrix^6^, contains a compact α/β domain that is structurally analogous to the isolated N-terminal domain in yeast MRS2^38^. It is followed by an α-helical domain that extends into the inner membrane via an extensively elongated α-helix, the ‘stalk’ helix (Fig. 1c, Extended Data Fig. 3a). This stalk helix bridges the N-terminal matrix domain and the transmembrane pore by forming a continuous α-helix with the first transmembrane helix TM1, which lines the central ion pathway (Fig. 1c). Following the inner helix TM1, an ordered short loop, containing the GMN motif, reaches into the intermembrane space and constitutes the ion entryway. The outer helix TM2 mainly packs against TM1 from the same subunit in the pentameric channel, with its C-terminus residing in the mitochondrial matrix (Fig. 1c). Despite low sequence similarity, the overall architecture of human MRS2 is reminiscent of CorA and ZntB^18–21,32,39^, with each subunit adopting a similar fold (Extended Data Fig. 3).

**Fig. 1.**
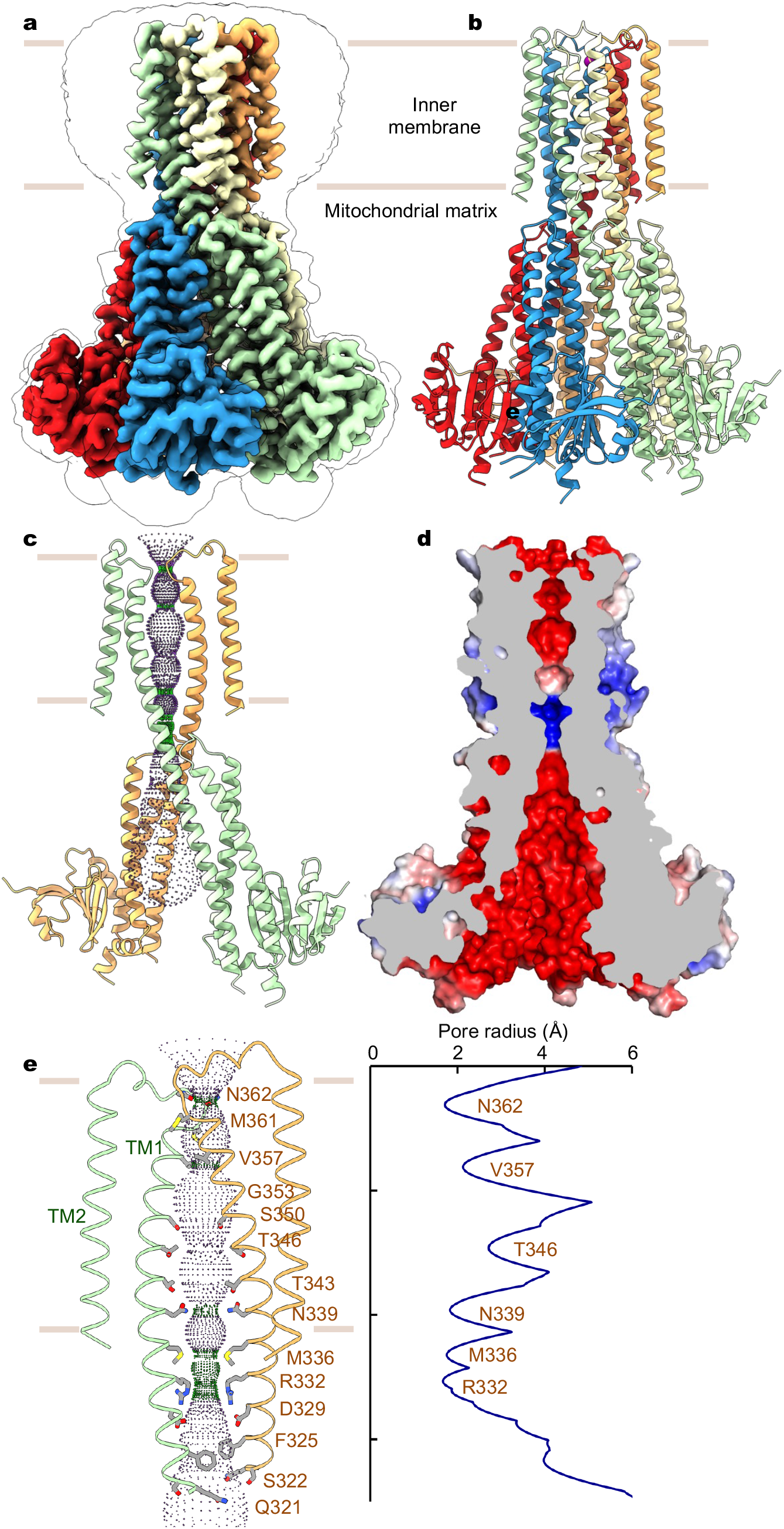
Structure of human MRS2. **a**, Cryo-EM reconstruction of human MRS2. Each subunit is uniquely colored and the contour indicates membrane boundaries. **b**, The overall structure of human MRS2. **c**, The central ion pathway, estimated using the HOLE program and represented as colored dots (pore radius: 1.15 Å < green < 2.3 Å < blue). Two opposite subunits are shown for clarity. **d**, Cutaway view of the channel surface, colored by surface electrostatic potential (red, −5 kT/e; white, neutral; blue, +5 kT/e). **e**, Details of the pore-lining residues and the dimension of the central pore.

However, while the transmembrane domain (TM1-TM2) and the α-helical domain are highly conserved among these proteins, the peripheral α/β domain in MRS2 is smaller in size than that of CorA (∼90 vs. ∼150 amino acids), thus resulting in different topology and structural arrangement. In particular, the α/β domain in human MRS2 contains a six-stranded β-sheet with two α-helices, instead of a seven-stranded β-sheet with four α-helices as observed in that of CorA (Extended Data Fig. 3). In CorA, Mg^2+^ occupancy at the α/β domain inter-subunit interface dictates the distribution of distinct conformations with varying degrees of symmetry break of the pentameric channel and confers Mg^2+^-dependent closure of the transmembrane ion pore^24,25^. Thus, the contrasting α/β domain structure and assembly interface may hint at a distinct gating mechanism of MRS2 from CorA (Extended Data Fig. 3).

### Ion pathway and binding sites

The ion conduction path of MRS2 is primarily defined by the inner helix TM1, which spans the inner membrane and extends into the matrix (Fig. 1c). The central ion pore, predominantly lined by polar amino acids, is largely electronegative (Fig. 1d, e), thus supporting permeation of cations. Notably, at the membrane-matrix interface, two narrow constriction sites are generated by a ring of methionine (M336) and a ring of arginine (R332) residues, respectively. In particular, the electropositive arginine ring, highly conserved in mammalian MRS2 proteins but absent in the prokaryotic CorA channels (Extended Data Fig. 4), would present an energy barrier for cation movement. This energy barrier, however, is partially compensated by the electronegative aspartate ring one helical turn below. Beyond this narrow region, the vestibule in the matrix widens, and its electronegative surface is compatible with cation conduction (Fig. 1d, 1e).

In the divalent-free MRS2_EDTA_ structure, discernible ion densities are absent in the central pore. By contrast, the MRS2_Mg_ and MRS2_Ca_ structures show robust densities consistent with these respective ions (Fig. 2, Extended Data Fig. 5). Alignment of these three structures, MRS2_EDTA_, MRS2_Mg_ and MRS2_Ca_, reveals that the channel maintains a static structure, and notable differences are only present at the ion binding sites in the central pore and at the inter-subunit assembly interfaces in the matrix (Fig. 2).

**Fig. 2.**
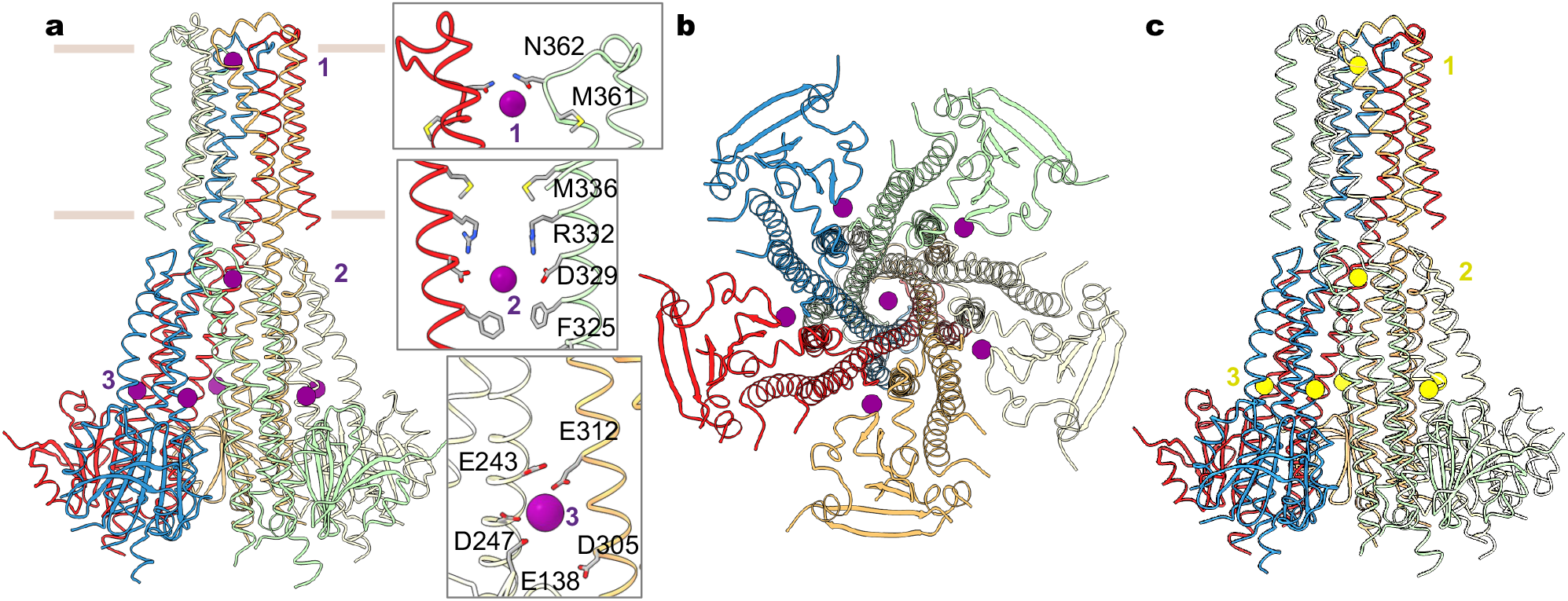
Mg^2+^ and Ca^2+^ recognition. **a**, Mg^2+^ ions (purple spheres) identified in the structure of MRS2 in the presence of Mg^2+^. Also shown are the details of the three Mg^2+^ ions (site 1 and 2) coordinated within the channel. Only two adjacent subunits are shown for clarity. **b**, Orthogonal view as in (**a**). **c**, Ca^2+^ ions (yellow spheres) in the MRS2_Ca_ structure.

Divalent ions, Mg^2+^ and Ca^2+^, are similarly bound to the channel. In the central pore, two distinct ion binding sites are generated by the pore-lining residues. Site 1, located at the entrance of the intermembrane space, is constituted by N362 of the GMN motif, together with the backbone carbonyl oxygen atoms of G360 (Fig. 2a). The equivalent site in the prokaryotic CorA channels was suggested to form a selectivity filter^22,40,41^. Site 2 is near the acidic D329 ring in the mitochondrial matrix. Of note, an analogous site 2 is present in CorA, but the location of this ion binding site is shifted one helical turn further into the cytoplasmic side^41^ (Extended Data Figs. 3, 4). The corresponding residue of D329 in TmCorA is serine, and the acidic aspartate ring (D277 in TmCorA) coordinating Mg^2+^ is one helical turn away (Extended Data Figs. 3, 4). An additional unique Mg^2+^ binding site (site 3) is generated by an acidic pocket at the inter-subunit interface, which is constituted by E138, E243 and D247 from one subunit and E312 from an adjacent subunit (Fig. 2). Notably, this interfacial binding site differs from those in TmCorA, which are critical for Mg^2+^-dependent inhibition of channel activity (Extended Data Fig. 3). It appears that the evolved interfacial Mg^2+^-binding site in MRS2, together with reduced inter-subunit packing interactions owing to a smaller α/β domain, suggests a likely distinct channel regulation mechanism. Furthermore, the similar divalent ion recognition sites observed in the MRS2_Mg_ and MRS2_Ca_ structures suggest that MRS2 may conduct both Mg^2+^ and Ca^2+^.

### Expression and recordings in *Xenopus* oocytes

To investigate the functional properties, we expressed the full-length human MRS2 protein (MRS2_FL_) or MRS2_EM_ heterologously in *Xenopus* oocytes, which have been used for electrophysiological recordings of TmCorA^22,23^. Consistent with previous reports, two-electrode voltage clamp (TEVC) showed that perfusion of TmCorA-expressing oocytes with a Mg^2+^-containing solution elicited large inward currents that peaked within seconds and decayed slowly following a course of several minutes (Fig. 3a). However, the same procedure failed to induce Mg^2+^ currents for MRS2_FL_ or MRS2_EM_ (Fig. 3a), although both exhibited higher levels of surface expression than TmCorA (Fig. 3a).

**Fig. 3.**
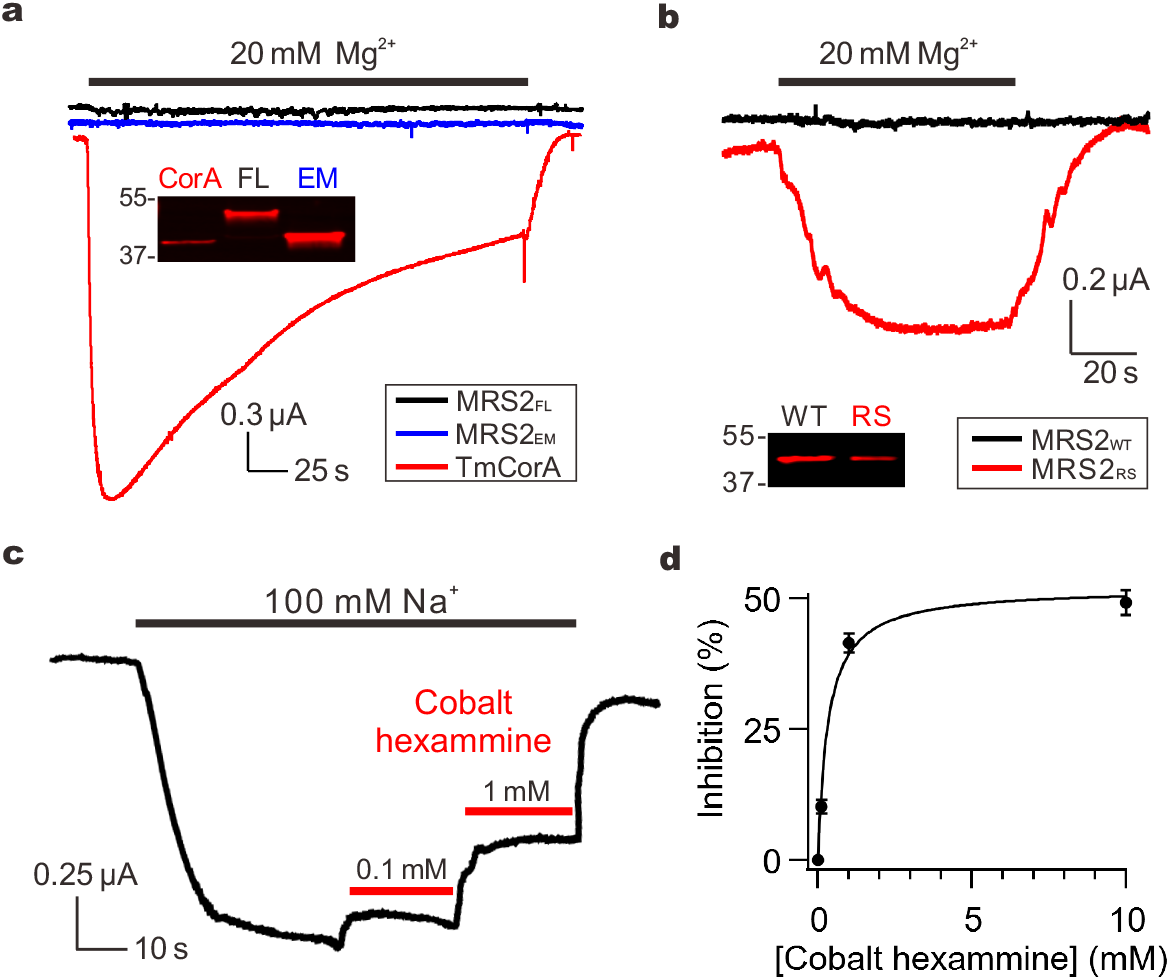
MRS2 currents. **a**, Mg^2+^ transport by TmCorA or MRS2. The western blot image compares the expression of TmCorA, MRS2_FL_, and MRS_EM_, all tagged with a C-terminal 1D4 sequence and detected with an anti-1D4 antibody. **b,** MRS2 Mg^2+^ currents induced by the R332S mutation. MRS2_RS_ exhibits ∼60% expression level of MRS2_WT_. **c,** Inhibition of MRS2 activity by cobalt hexammine. Na^+^ currents were presented here, but similar inhibition was obtained when recording Mg^2+^ currents. **d,** Dose response of cobalt hexammine inhibition of MRS2 Na^+^ currents. Data points were fit with a standard saturation binding equation (black curve). Maximal inhibition achieved is ∼50%, comparable with that obtained with TmCorA.

Structural analysis of MRS2 suggests that the narrow, electropositive R332 ring (Fig. 1e) might present an energy barrier that suppresses cation permeation. Supporting this notion, replacement of R332 with a smaller polar amino acid, serine (R332S), produced µA levels of Mg^2+^ currents (Fig. 3b). Thus, charge neutralization and reduced size of the side chain at this constriction site greatly facilitated cation conduction. We reasoned that these observed activities were mediated by MRS2, because (1) they were not seen in oocytes without MRS2 expression or oocytes expressing the mitochondrial Ca^2+^ uniporter, and (2) they could be reduced by a known CorA inhibitor cobalt hexammine (Fig. 3c), with an IC_50_ of 0.3 mM (Fig. 3d). Of note, when we were preparing this manuscript, an independent structural study of human MRS2 was published and suggested that R332 might bind Cl^-^ to facilitate Mg^2+^ transport^42^. This Cl^-^-mediated Mg^2+^ transport mechanism is not compatible with our results of R332S, which would abolish Cl^-^ binding but drastically enhanced Mg^2+^ permeation. Moreover, Mg^2+^ currents were not observed in wild-type (WT) MRS2 in a recording solution with >100 mM of Cl^-^ (Fig. 3a). Lastly, we consistently obtained larger Mg^2+^ currents with the R332S substitution in MRS2_EM_ than MRS2_FL_, likely because the former displayed higher surface expression (Fig. 3a). This observation suggests that the MRS2_EM_ construct is an appropriate surrogate for MRS2_FL_ in our structural and functional analysis. Moreover, the more robust activity of MRS2_EM_ allows detailed electrophysiological characterization; we thus perform functional studies on the MRS2_EM_ background in this study. For clarity, hereinafter we refer WT MRS2_EM_ and the R332S mutant of MRS2_EM_ as MRS2_WT_ and MRS2_RS_, respectively.

### Ion permeation and channel regulation

Unlike TmCorA (Fig. 3a), MRS2 showed no sign of Mg^2+^ inactivation (Fig. 3b), corroborating the notion that MRS2 might have evolved unique functional properties. This motivated us to further characterize ion permeation and regulation of MRS2. TEVC experiments varying cations in the perfusion solutions showed that MRS2_RS_ could conduct Ca^2+^, K^+^ and Na^+^ (Fig. 4a), and the currents for all these ions are drastically diminished without the R332S mutation (Fig. 4a).

**Fig. 4.**
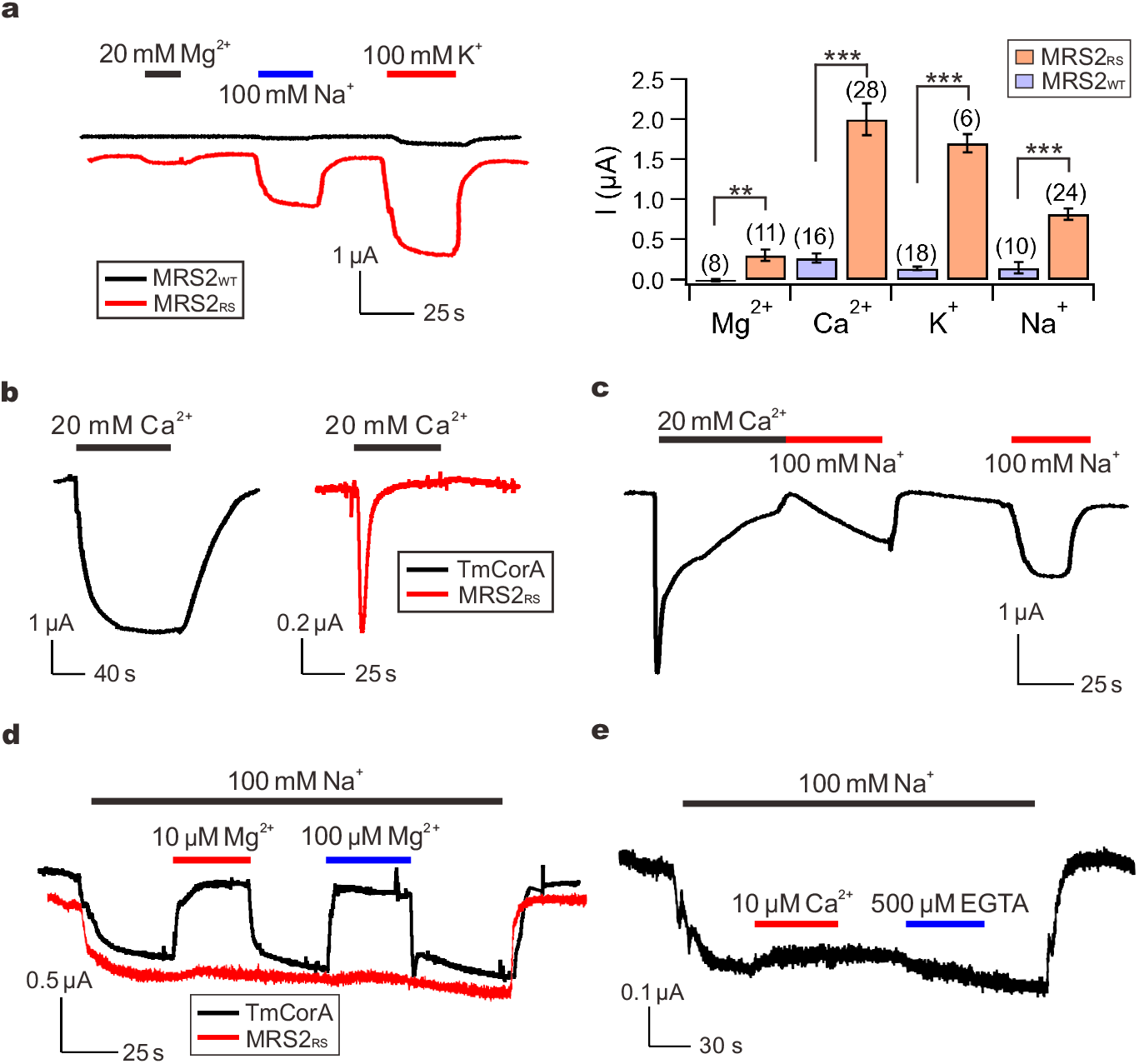
Ion permeation properties. **a**, Monovalent and divalent cation currents through MRS2_WT_ or MRS2_RS_. Peak-level amplitudes of Ca^2+^ currents were presented in the bar chart. **P < 0.01; ***p < 0.001. **b,** Ca^2+^ currents conducted by TmCorA or MRS2_RS_. **c,** Recovery of MRS2_RS_ from inactivation. **d,** The effect of Mg^2+^ on Na^+^ currents. **e,** Lack of Ca^2+^ inhibition of MRS2 Na^+^ currents. 500 µM of EGTA was added later in the recording to test if the trace amount of Ca^2+^ in the NaCl buffer can inhibit Na^+^ currents.

Interestingly, the Ca^2+^ currents, but not K^+^ or Na^+^ currents, showed a rapid decay to a steady state of less than 5% of the peak level within 1 minute (Fig. 4a-b). We concluded that these inactivating Ca^2+^ currents were not caused by contaminating Ca^2+^-activated Cl^-^ currents (CACC)^43–45^ for three reasons. First, no such inactivation was observed in our recordings of TmCorA, which is known to conduct Ca^2+^ ^22^(Fig. 4b). Second, these Ca^2+^ currents could still be observed with injection of 5 nmol of BAPTA into oocytes to suppress CACC. Finally, when oocytes were acutely exposed to a Ca^2+^-free, Na^+^-containing solution following the decay of Ca^2+^ currents, the Na^+^ currents increased slowly with a time course of tens of seconds, suggesting that MRS2_RS_ was inactivated (Fig. 4c). By contrast, if the Na^+^-containing solution was applied several minutes after Ca^2+^ withdrawal to allow the channels to recover from the inactivated state, Na^+^ currents would reach peak levels within seconds (Fig. 4c). Altogether, these results suggest that physiologically MRS2 might be regulated by matrix Ca^2+^ instead of Mg^2+^.

It has been reported that CorA, also known to permeate K^+^ and Na^+^, can selectively conduct Mg^2+^ by a classical multi-ion pore repulsion mechanism^22^. Specifically, in the presence of submicromolar [Mg^2+^], Mg^2+^ binding to a high-affinity site within the pore would block the passage of incoming monovalent cations. When [Mg^2+^] rises to the physiological range of millimolar levels, additional Mg^2+^ entering the pore would create electrical repulsion between these Mg^2+^ ions, thus leading to rapid Mg^2+^ permeation. This phenomenon is known as the anomalous mole fraction effect (AMFE), which is also recapitulated in our experimental system (Fig. 4d). However, MRS2_RS_-mediated Na^+^ currents (100 mM Na^+^) are not affected by submicromolar levels of Mg^2+^ or Ca^2+^, suggesting that MRS2 does not selectively transport Mg^2+^ or Ca^2+^ (Fig. 4d-e). Taken together, we conclude that MRS2 is a Ca^2+^-regulated nonselective cation channel.

### Functional roles of divalent cation sites

The polar asparagine ring from the GMN motif, conserved in the CorA family of Mg^2+^ channels, generates a divalent cation binding site (site 1 in Fig. 2a), presumably playing a role in ion selectivity. Consistent with this notion, it has been shown that alanine substitution (N314A) compromises, but not fully eliminates, Mg^2+^ blockade of Na^+^ currents in TmCorA^22^. However, the detailed atomic basis of ion selectivity remains elusive as the asparagine ring in the CorA family proteins has been shown to be compatible with binding of multiple divalent ions including Mg^2+^, Ni^2+^, Co^2+^ and Zn^2+^ ^32,39^. Notably, this GMN cation binding site is not necessary for ion permeation, because µA levels of whole-oocyte currents can still be obtained with the TmCorA N314A mutant^22^. The lack of AMFE and Mg^2+^ selectivity in MRS2 raises the possibility that its GMN cation site, site 1, might play a limited role in channel function. Indeed, we find that the N362A substitution in the GMN motif in MRS2_RS_ only reduces Na^+^ or Ca^2+^ currents by ∼50% (Fig. 5a-b).

**Fig. 5.**
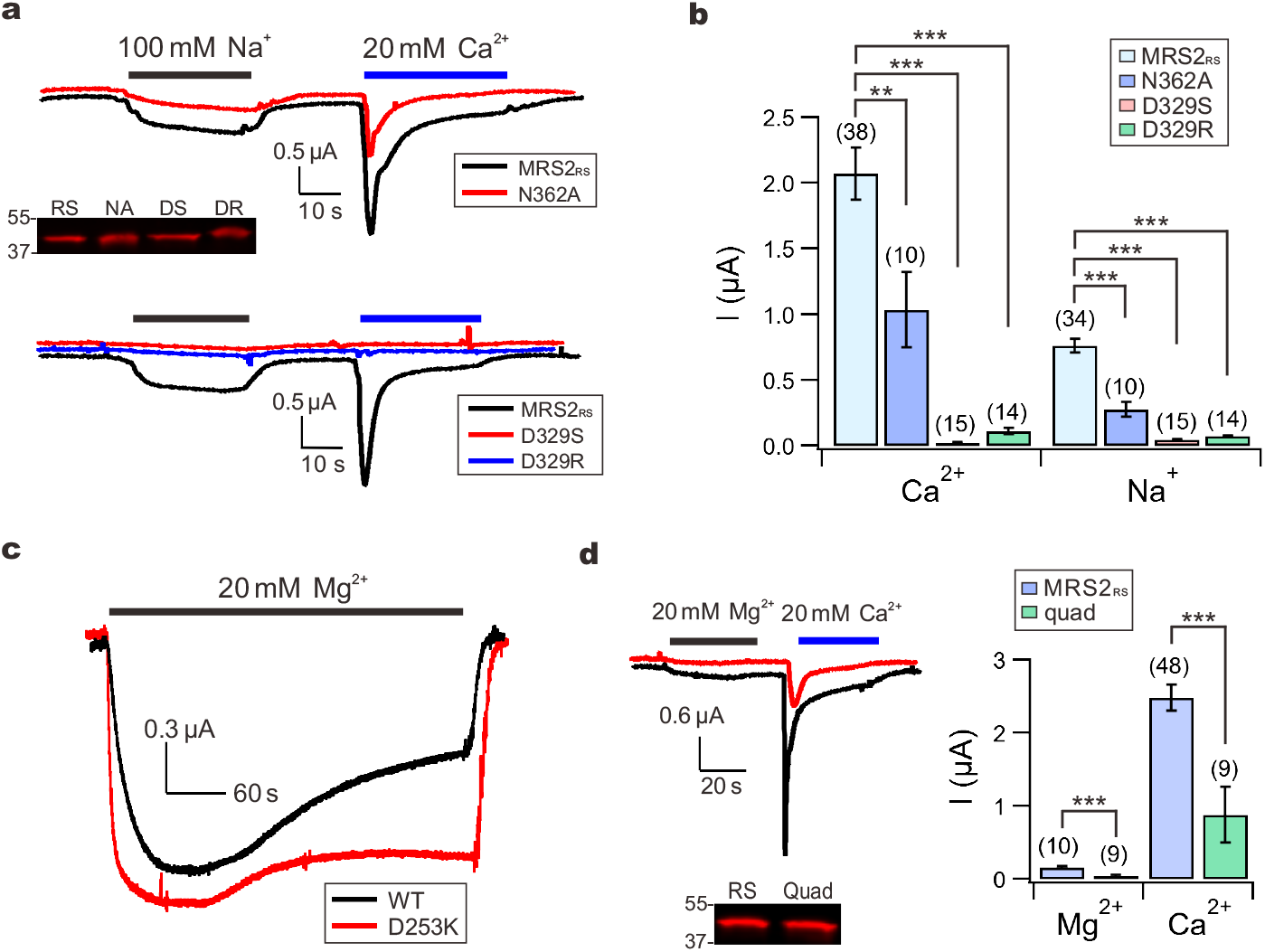
Divalent cation-binding sites. **a-b**, Functional impacts of site 1 or site 2 mutations on MRS2_RS_. The Western-blot image shows that these mutations do not affect MRS2_RS_ expression. *RS*: MRS2_RS_. *NA*: N362A; *DS*: D329S; *DR*: D329R. **c**, Mg^2+^ inactivation of TmCorA. **d,** The effect of site 3 mutations on MRS2_RS_ function (q*uad*: E138A-E243A-D247A-E312A mutation). Western blot shows that these mutations do not reduce surface expression.

In our MRS2 structures, a second divalent cation binding site (Fig. 2a, Extended Data Fig. 3) is located on the matrix end in the central pore, formed by a highly conserved aspartate residue, D329, one helical turn away from R332. Serine or arginine substitution of D329 in MRS2_RS_ drastically reduces Na^+^ or Ca^2+^ currents (Fig. 5a-b), indicating that the D329 cation site, site 2, is critical for ion permeation in MRS2.

The inter-subunit Mg^2+^ binding sites found in the cytoplasmic domain of TmCorA have been shown to mediate Mg^2+^ inactivation^22–24^. Consistent with previous reports, we find that disruption of this Mg^2+^ site by a charge-reversal mutation (D253K) in TmCorA strongly suppresses, but does not entirely abolish, Mg^2+^ inactivation (Fig. 5c). Our MRS2 structures reveal a distinct inter-subunit divalent cation binding site, site 3, that is much closer to the transmembrane region (Fig. 2, Extended Data Fig. 3) and is formed by a group of acidic residues (E138, E243, D247, and E312). Given the regulatory role of the cytoplasmic Mg^2+^ site in TmCorA, we speculated that site 3 might be responsible for Ca^2+^ inactivation in MRS2. Surprisingly, an E138A-E243A-D247A-E312A MRS2_RS_ mutant maintained robust Ca^2+^ inactivation, although the current levels were ∼70% smaller (Fig. 5d). It thus appears that site 3 contributes to ion conduction without a direct role in Ca^2+^ regulation.

## Discussion

This work now defines the structural and functional properties of human MRS2 channel, decades after its molecular identification and proposed role as a Mg^2+^ transporter. These findings greatly advance our understanding of mitochondrial Mg^2+^ transport and address a longstanding conundrum in the field. Accumulating evidence suggests that MRS2 forms the mitochondrial Mg^2+^ entryway. However, why is free [Mg^2+^] in the matrix only comparable to that in the cytosol, ∼0.3-1 mM^46–49^? In principle, Mg^2+^ channels should be able to support a 10^6^-fold, cytosol-directed Mg^2+^ gradient with a typical −180 mV inner membrane potential. It turns out that MRS2 possesses an unusual molecular property in that its ion permeation is energetically impeded by a conserved basic arginine ring within the pore, which is absent in other CorA-family proteins. Consequently, Mg^2+^ influx can be more readily countered by the slow mitochondrial Mg^2+^ efflux mechanisms mediated by Mg^2+^ transporters^29,50,51^, thus leading to comparable [Mg^2+^] inside and outside of the matrix. We speculate that the arginine ring, which slows ion conduction in MRS2, is evolved out of physiological necessity. With low millimolar levels of free Mg^2+^ in the cytosol, a typical ion channel would likely conduct Mg^2+^ into the matrix quickly to collapse the inner membrane potential, thus diminishing the driving force for mitochondrial ATP synthesis. Indeed, it is well established that ionophores, such as Carbonyl cyanide-*p*-trifluoromethoxyphenylhydrazone (FCCP), can readily depolarize mitochondrial membranes even with much lower turnover rates than typical ion channels^52,53^. Of note, the limited MRS2 ion conduction rate also explains earlier findings of slow ^28^Mg influx into mitochondria and a lack of Mg^2+^ currents across the inner membrane in the presence of 1 mM extra-mitochondrial Mg^2+^ and absence of Ca^2+^ from mitoplast patch-clamp recordings^54^.

Our discovery of MRS2 functioning as a Ca^2+^-regulated nonselective cation channel, permeable to not only Mg^2+^ but also Ca^2+^, K^+^ and Na^+^, may have profound implications in mitochondrial ion homeostasis. It has been established for decades that mitochondria operate Na^+^ and K^+^ cycles to control mitochondrial Na^+^ and K^+^ homeostasis. Specifically, Na^+^ and K^+^ are imported into the matrix via slow, electrophoretic Na^+^ and K^+^ uniport mechanisms, and are extruded via Na^+^/H^+^ and K^+^/H^+^ exchangers, respectively^31,55,56^. Na^+^ is known to regulate mitochondrial Ca^2+^-dependent processes, such as oxidative phosphorylation and cell death, by participating in Na^+^/Ca^2+^ antiport^57–59^, while K^+^ plays a prominent role in regulating mitochondrial volume and inner membrane potential^31,60^. To date, multiple protein components have been proposed to mediate mitochondrial Na^+^ or K^+^ transport, but consensus is lacking regarding the exact function, localization, and activation conditions of these proteins^61,62^. We reason that MRS2 may be well positioned to contribute to mitochondrial Na^+^ and K^+^ uniport in that its slow kinetics could mitigate the problem of mitochondrial inner-membrane potential dissipated by typical channels with fast kinetics.

In contrast to CorA, which is inhibited by cytoplasmic Mg^2+^, MRS2 is not regulated by matrix Mg^2+^ but instead inactivated by Ca^2+^. It has been shown that the Mg^2+^ inactivation process of CorA requires fast Mg^2+^ influx to significantly alter local [Mg^2+^] near the cytoplasmic exit of the channel, as demonstrated by more rapid channel inactivation induced by faster permeant Co^2+22^. Since Mg^2+^ permeation via MRS2 is at least 50-fold slower than that in CorA, it is unlikely that such slow Mg^2+^ flux can drastically alter the millimolar levels of Mg^2+^ present in the matrix to exert efficient regulation. By contrast, during eukaryotic evolution, Ca^2+^ becomes a critical secondary messenger in multicellular organisms, which keep cytoplasmic Ca^2+^ at submicromolar levels so that opening of Ca^2+^ channels can effectively elevate local [Ca^2+^] to initiate downstream Ca^2+^-dependent processes^63^. Perhaps MRS2 evolves from the prokaryotic Mg^2+^-regulated CorA to become a Ca^2+^-regulated channel to adapt to the more prominent role of Ca^2+^ in metazoan physiology. We note that under physiological conditions, Ca^2+^ permeation via MRS2 would be negligible because cytosolic [Ca^2+^] is >1000-fold lower than [Mg^2+^], [Na^+^], and [K^+^]. Therefore, the observed Ca^2+^ inactivation of MRS2 unlikely reflects a negative feedback process occurred in voltage-gated Ca^2+^ channels, wherein Ca^2+^ influx shuts the channel to terminate the Ca^2+^ flux itself. Instead, MRS2 might serve as a component in the mitochondrial Ca^2+^ signaling pathways, modulating mitochondrial metabolism in response to fluctuations in matrix [Ca^2+^] mediated by mitochondrial Ca^2+^ transport proteins.

Surprisingly, disruption of Ca^2+^ binding at site 3 did not abolish Ca^2+^ regulation of MRS2. However, this is indeed consistent with the structural observation that site 3 can bind not only Ca^2+^ but also Mg^2+^. The essentially identical MRS2 structures in different ionic conditions, including Na^+^, Mg^2+^ and Ca^2+^, suggest a subtle regulation mechanism by matrix Ca^2+^ that remains to be elucidated. The rather rigid structure of MRS2 contrasts with that of CorA, in which an ensemble of conformations, including apparently asymmetric assemblies of the pentameric channel, exists in the absence of Mg^2+^. Mg^2+^ binding at the intracellular inter-subunit interface in CorA reduces structural dynamics and promotes a symmetric, closed conformation^24,25^. In contrast, the pronounced asymmetry or conformational heterogeneity is not observed in human MRS2 in the absence or presence of divalent cations. These distinct features further support the evolved structural and functional properties of MRS2, which now provide a foundation to further examine the physiological roles of MRS2 in mitochondrial biology.

## Acknowledgements

This work was partly supported by NIH grants R01NS109307 (to P.Y.) and R01GM144485 (to M.T.). We thank Tsung-Yun Liu for technical assistance. We thank the staff scientists at Washington University Center for Cellular Imaging for data acquisition. Some of this work was performed at the Simons Electron Microscopy Center at the New York Structural Biology Center, with major support from the Simons Foundation (SF349247).

## Author Contributions

Z.H. performed protein biochemistry. Z.H., J.M. and J.Z. collected cryo-EM images. J.M., J.Z. and P.Y. conducted cryo-EM structure determination and analysis. Y.T. and C.T. performed electrophysiology experiments. M.T. and P.Y. supervised the project and wrote the manuscript. All authors edited the manuscript.

## Methods

### Cloning, expression and purification

The codon-optimized DNA fragment encoding *H. sapiens* MRS2 (hMRS2, NCBI sequence: NP_065713.1) was synthesized (Bio Basic Inc.) and cloned into a modified yeast *P. pastoris* expression vector pPICZ-B with a PreScission protease cleavage site followed by a C-terminal GFP-His_10_ tag. Initial FSEC screen using the full-length wild-type protein showed very low expression level. The N-terminal 61 amino acids corresponding to the mitochondrial targeting sequence were removed from the expression construct. Additionally, removal of the C-terminal 12 amino acids, which are predicted to be unstructured, improved the protein expression level but was insufficient for structural studies. Fusion of BRIL (thermostabilized apocytochrome b562RIL) further improved expression and biochemical stability. The final expression construct, MRS2_EM_, contains residues 62 to 431 of human MRS2 and a C-terminal BRIL followed by the PreScission protease cleavage site and GFP-His_10_ tag.

For electrophysiological recordings, TmCorA and MRS2 constructs were fused with a C-terminal 1D4 tag (TETSQVAPA), and cloned into a pOX vector^73^. Site-directed mutagenesis was performed using standard two-step PCR mutagenesis using the Herculase II DNA Polymerase (Agilent), with sequences verified by Sanger sequencing. For *in vitro* transcription, the pOX vector was linearized with NotI, wih 10 µg of the linearized vector used for complementary RNA (cRNA) synthesis using the mMESSAGE mMACHINE T3 transcription kit (Thermo Fisher). The cRNA was purified using the RNeasy MinElute Cleanup kit (Qiagen), dissolved in nuclease-free water to a final concentration of ∼1.8 µg/µl, and then stored at −80 °C. The quality of cRNA was verified using electrophoresis with 4% formaldehyde to denature the cRNA.

Yeast cells expressing MRS2_EM_ were disrupted by mechanical milling (Retsch MM400) and resuspended in buffer containing 50 mM Tris pH 8.0 and 150 mM NaCl in the presence of protease inhibitors (2.5 μg ml^-1^ leupeptin (L-010-100, GoldBio), 1 μg ml^-1^ pepstatin A (P-020-100, GoldBio), 100 μg ml^−1^ 4-(2-Aminoethyl) benzenesulfonyl fluoride hydrochloride (A-540-10, GoldBio), 3 μg ml^−1^ aprotinin (A-655-100, GoldBio), 1 mM benzamidine (B-050-100, GoldBio), 200 μM phenylmethane sulphonylfluoride (P-470-25, GoldBio) and DNase I (D-300-1, GoldBio). The cell mixture was extracted with 1% (w/v) lauryl maltose neopentyl glycol (LMNG, NG310, Anatrace) for 2 hours with stirring at 4℃ and then centrifuged for 1 hour at 30,000g. The supernatant was collected and incubated with 3 ml of cobalt-charged resin (786-403, G-Biosciences) for 3 hours at 4℃. Resin was then collected and washed with 10 column volumes of buffer containing 20 mM Tris pH 8.0, 150 mM NaCl, 20 mM imidazole and 85 μM glyco-diosgenin (GDN, GDN101, Anatrace). The protein was eluted with 200 mM imidazole and digested with PreScission protease at 4 ℃ overnight to remove the C-terminal GFP-His_10_ tag. The digested protein was concentrated using a 100 kDa concentrator and further purified on a Superose 6 Increase 10/300 gel filtration column (GE Healthcare Life Sciences) equilibrated in an Mg^2+^-containing buffer (20 mM Tris pH 8.0, 150 mM NaCl, 40 mM MgCl_2_ and 40 μM GDN) or EDTA buffer (20 mM Tris pH 8.0, 150 mM NaCl, 10 mM EDTA and 40 μM GDN) or Ca^2+^-containing buffer (20 mM Tris pH 8.0, 150 mM NaCl, 10 mM Ca^2+^ and 40 μM GDN). Peak fractions containing channel protein were concentrated to ∼5 mg ml^-1^ and used immediately for cryo-EM grid preparations.

### Cryo-EM grid preparation and imaging

A volume of 3.5 μl of purified channel protein was applied onto glow-discharged copper holey carbon grids R1.2/1.3 (Q350CR1.3, Electron Microscopy Sciences), which was cleaned for 1 minute in an H_2_/O_2_ plasma using a Gatan Solarus 950 (Gatan). Grids for MRS2_EDTA_ and MRS2_Mg_ were blotted for 2 s at ∼100% humidity, and grids for MRS2_Ca_ blotted at 80% humidity with a blot force of 0. Grids were then plunge frozen in liquid ethane using a Vitrobot Mark IV (ThermoFisher Scientific). The MRS2_EDTA_ and MRS2_Mg_ grids were loaded onto a Glacios (FEI) electron microscope operating at an accelerating voltage of 200 kV with Falcon 4 Detector (ThermoFisher Scientific). Images were recorded with EPU software (ThermoFisher Scientific) in counting mode with a pixel size of 1.2 Å and a nominal defocus value ranging from −1.0 to −2.4 μm. For the Mg^2+^-bound sample, single-particle cryo-EM data were collected with a dose of ∼5.56 electrons per Å^2^ per second, and each movie was recorded with a 7.33 s exposure with 42 total frames (representing an accumulated dose of ∼41 electrons per Å^2^). For the EDTA sample, images were collected with a dose of ∼6.66 electrons per Å^2^ per second, and each movie was recorded with a 7.33 s exposure with 42 total frames (representing an accumulated dose of ∼49 electrons per Å^2^). The MRS2_Ca_ grids were loaded onto a Krios electron microscope operating at 300 kV equipped with a K3 detector (Gatan). Movies were recorded using the Leginon software^64^ with a nominal pixel size of 1.076 Å and defocus value between −0.8 to −2.5 μm. Data were collected with a dose rate of 25.89 electrons per Å^2^ per second over 40 frames and 2 s for a total dose of 51.77 electrons per Å^2^.

### Image processing and map calculation

Image processing and map calculation were performed in CryoSPARC^65^. Recorded movies were aligned and dose weighted and then subjected to contrast transfer function (CTF) estimation. Low-quality images were removed for further processing. Auto-picked particles were extracted and used to generate 2D classes, and classes representing an intact channel were selected for template-based particle picking. Several rounds of 2D classification were conducted to guide the selection of particles for 3D map calculation. Multiple *ab initio* models were then generated for heterogeneous refinement. Particles representing an intact channel were used for non-uniform refinement followed by local refinement. Maps calculated had overall resolutions in the range of ∼2.9 to 3.1 Å.

### Model building and coordinate refinement

The AlphaFold^33^ model of human MRS2 was used as an initial model to facilitate model building and subsequent adjustment in COOT^66^. Cycles of model building in COOT and real-space refinement using real_space_refine against the full maps in PHENIX^67^ were conducted. ISOLDE was used to improve model building^68^. The final refined atomic models were validated using MolProbity^69^. Pore radius calculation was performed using the program HOLE^70^. Structural figures were generated using PyMOL (pymol.org) and UCSF ChimeraX^71^. All software packages used for data processing and structural analysis except for cryoSPARC were accessed through the SBGrid consortium^72^.

### Electrophysiological recordings

Preparation of *Xenopus* oocytes and TEVC recordings have been described in our previous study^74^. Briefly, oocytes were purchased from the Xenopus One company, and were digested with 1.25 mg/mL type II collagenase (Worthington) at room temperature in a solution containing 96 mM NaCl, 2 mM KCl, 2.5 mM MgCl_2_, 5 mM HEPES, pH 7.4-NaOH. Stage V–VI oocytes were selected and incubated in 18°C in a ND96 solution (96 mM NaCl, 2 mM KCl, 2 mM CaCl_2_, 0.5 mM MgCl_2_, 5 mM HEPES, pH 7.4-NaOH). Oocytes were injected with 40 ng of cRNA using a Nanojet II microinjector (Drummond), and were used for experiments 2 days after injection.

TEVC signals were measured using an Oocyte Clamp OC-725B system (Warner), filtered at 1 kHz, and sampled at 2 kHz. Voltage and current electrodes were filled with 3-M KCl with a resistance of 0.5 – 1 MΩ. Data acquisition and membrane voltage control were performed with a pCLAMP-10/Digidata-1322A system (Axon). All recordings were performed with the membrane potential clamped at −60 mV, and with oocytes initially perfused in an NMG buffer, containing 100 mM N-methyl-D-glucamine, 10 mM HEPES, pH 7.4-HCl. To test cation conductance, the perfusion solution was changed to buffers containing monovalent or divalent cations (MgCl_2_ buffer: 80 mM NMG, 20 mM MgCl_2_, 10 mM HEPES; CaCl_2_ buffer: 80 mM NMG, 20 mM CaCl_2_, 10 mM HEPES; NaCl buffer: 100 mM NaCl, 10 mM HEPES; KCl buffer: 100 mM KCl, 10 mM HEPES; pH 7.4 for all solutions). 100 μM of EDTA was added to the NaCl buffer to record Na^+^ current conducted by TmCorA. All recordings were performed under room temperature with at least 3 independent batches of oocytes.

### Membrane preparation and Western blots

Oocyte plasma membranes were isolated as described in our previous study^74^. In short, 12 oocytes were homogenized using a Dounce homogenizer with 1.2 mL of ice-cold homogenization buffer (HB), containing 80 mM sucrose, 20 mM Tris, 5 mM MgCl_2_, 5 mM NaH_2_PO_4_, 1 mM EDTA, pH 7.4. The lysate was spun down for 30 s at 10 g at 4 °C. After that, 1 mL of the supernatant was carefully removed and replaced with 1 mL of ice-cold HB, and the pellet was then resuspended and spun down at 10 g. The same procedure was repeated two more times, but with centrifugation at 20 g and then 40 g. After the 40-g spin, 1 mL of the supernatant was removed, and the pellet was resuspended in the remaining 200 µL of HB. It was then spun down for 20 min at 14,000 g at 4°C to pellet the membrane. The membrane pellet was resuspended in 30 µL of HB for Western analysis, denatured in an SDS loading buffer, and then loaded onto 4–20% gels for electrophoresis. The protein samples were transferred to PVDF membranes (LI-COR) using a TransBlot Turbo transfer system (Bio-Rad). Membranes were blocked in a Tris-buffered saline (TBS)-based intercept blocking buffer (LI-COR), and were then incubated with a home-made 1D4 primary antibody diluted to 0.1 µg/mL in TBST (TBS plus 0.075% Tween-20) at 4 °C overnight. After incubating with goat anti-mouse IRDye 680RD (LI-COR) secondary antibody diluted 1:15,000 in TBST in room temperature for 1 hour, the signals were captured using a LI-COR Odyssey CLx imager. Band intensities were quantified using the LI-COR Image Studio software (version 5.2).

### Data availability

The cryo-EM maps have been deposited to Electron Microscopy Data Bank with accession codes EMD-41587 (https://www.ebi.ac.uk/pdbe/entry/emdb/EMD-41587), EMD-41588 (https://www.ebi.ac.uk/pdbe/entry/emdb/EMD-41588), EMD-41589 (https://www.ebi.ac.uk/pdbe/entry/emdb/EMD-41589). Atomic coordinates have been deposited to the Protein Data Bank (PDB) with accession codes 8TS1 (https://www.rcsb.org/structure/8TS1), and 8TS2 (https://www.rcsb.org/structure/8TS2), 8TS3 (https://www.rcsb.org/structure/8TS3). Correspondence and requests for materials should be addressed to M.T. or P.Y.

---

**Extended Data Fig. 1.**
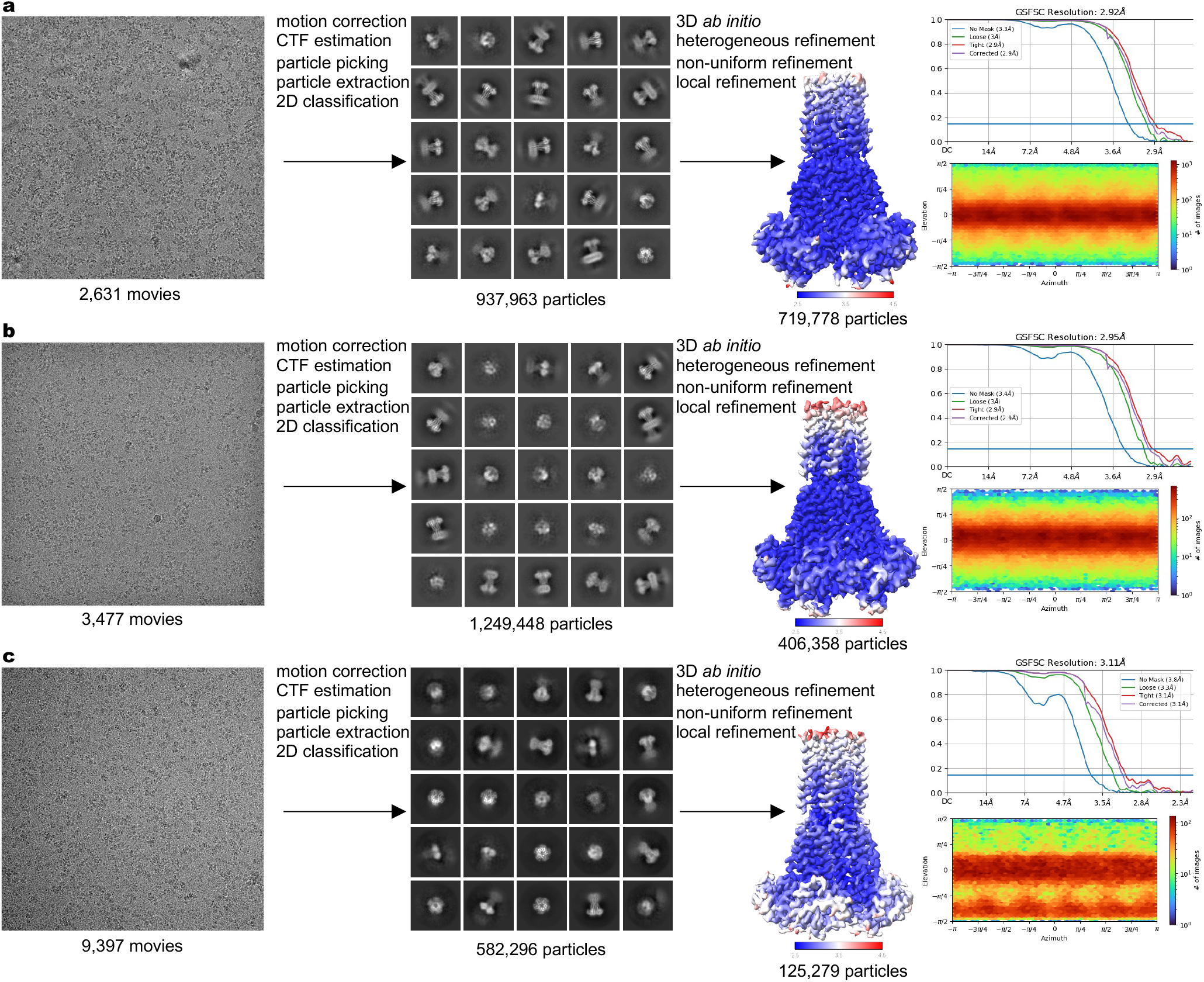
Cryo-EM data processing and validation. a-c, Image processing and map validation of MRS2Mg (a), MRS2EDTA (b), and MRS2Ca (c). Representative micrographs and 2D classes are shown. The final 3D reconstructions are colored by local resolution. Also shown are Fourier shell correlations (FSC) and orientation distribution of particles used in the final reconstruction.

**Extended Data Fig. 2.**
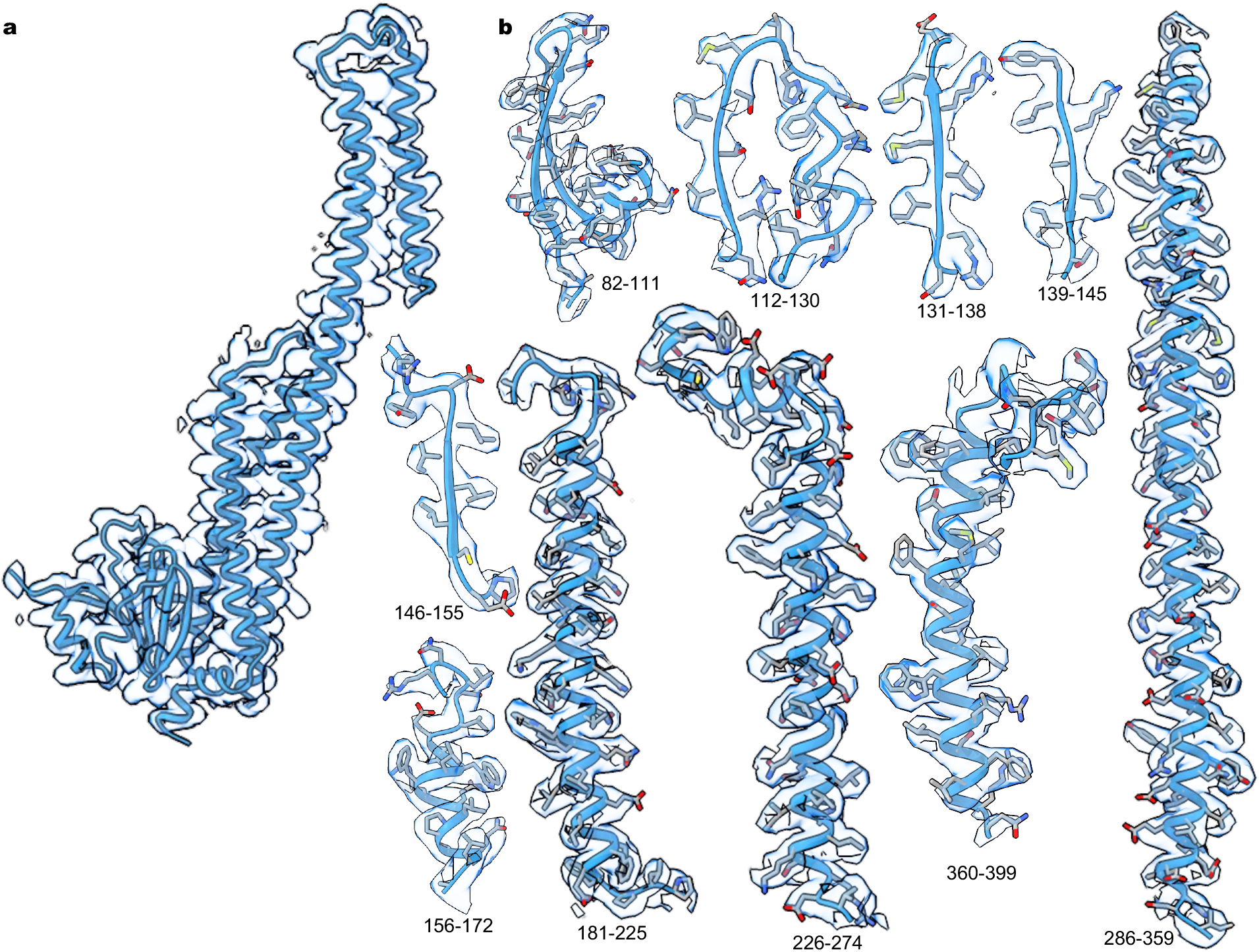
Cryo-EM density. a, Ribbon representation of a single subunit of MRS2Mg and cryo-EM density. b, Segments of the final refined model of MRS2Mg and the corresponding cryo-EM densities.

**Extended Data Fig. 3.**
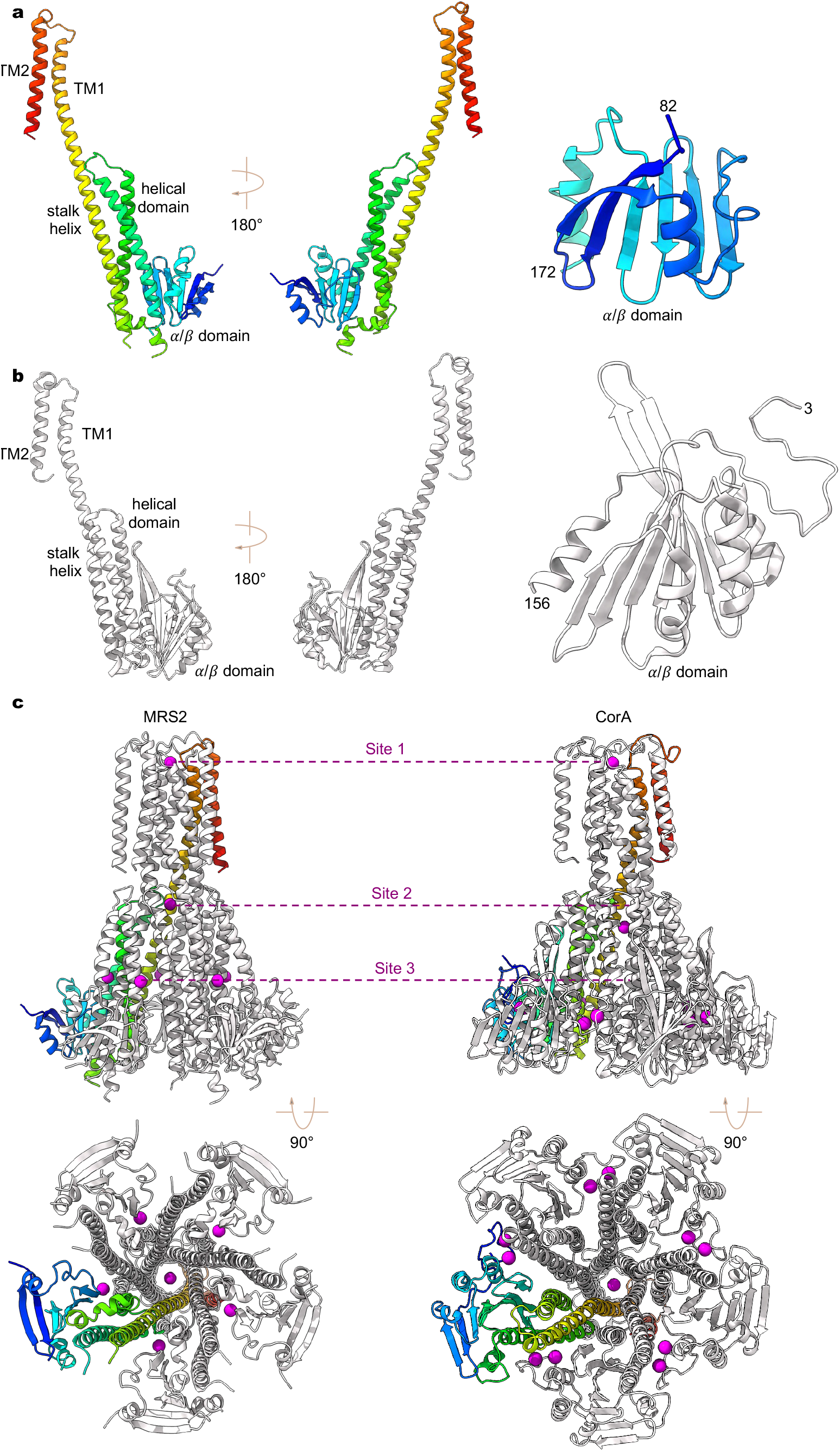
Structural comparison of human MRS2 and TmCorA. a-b, A single subunit of human MRS2 (a) and TmCorA (b, PDB ID: 4I0U). The N-terminal α/μ domain is also highlighted for comparison. c, Divalent ion binding sites in human MRS2 (left panels) and TmCorA (right panels).

**Extended Data Fig. 4.**
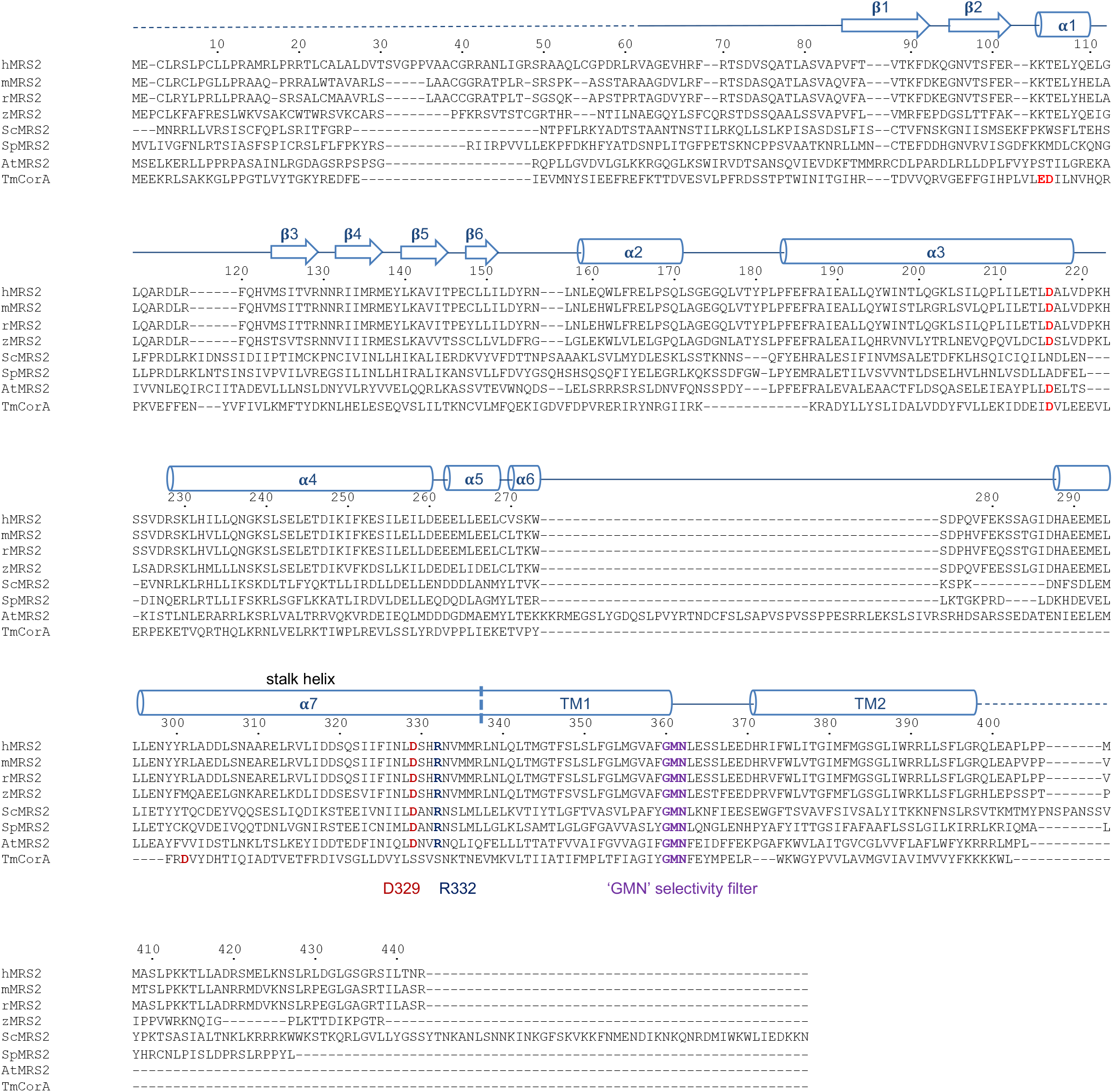
Sequence alignment of MRS2 channels. Multiple protein sequences, including *Homo sapiens* MRS2 (hMRS2, NCBI sequence: NP_065713.1), *Mus musculus* MRS2 (mMRS2, NCBI sequence: NP_001013407.2), *Rattus norvegicus* MRS2 (rMRS2, NCBI sequence: NP_076491.1), *Danio rerio* MRS2 (zMRS2, NCBI sequence: XP_693621.5), *Saccharomyces* cerevisiae MRS2 (ScMRS2, NCBI sequence: NP_014979.1), *Schizosaccharomyces pombe* MRS2 (SpMRS2, NCBI sequence: NP_596358.1), *Arabidopsis thaliana* MRS2 (AtMRS2, NCBI sequence: AAM62917.1), and *Thermotoga maritima CorA* (TmCorA, NCBI sequence: WP_004081315.1). Secondary structure elements on the basis of hMRS2 are indicated above the sequences. Critical amino acids are highlighted.

**Extended Data Fig. 5.**
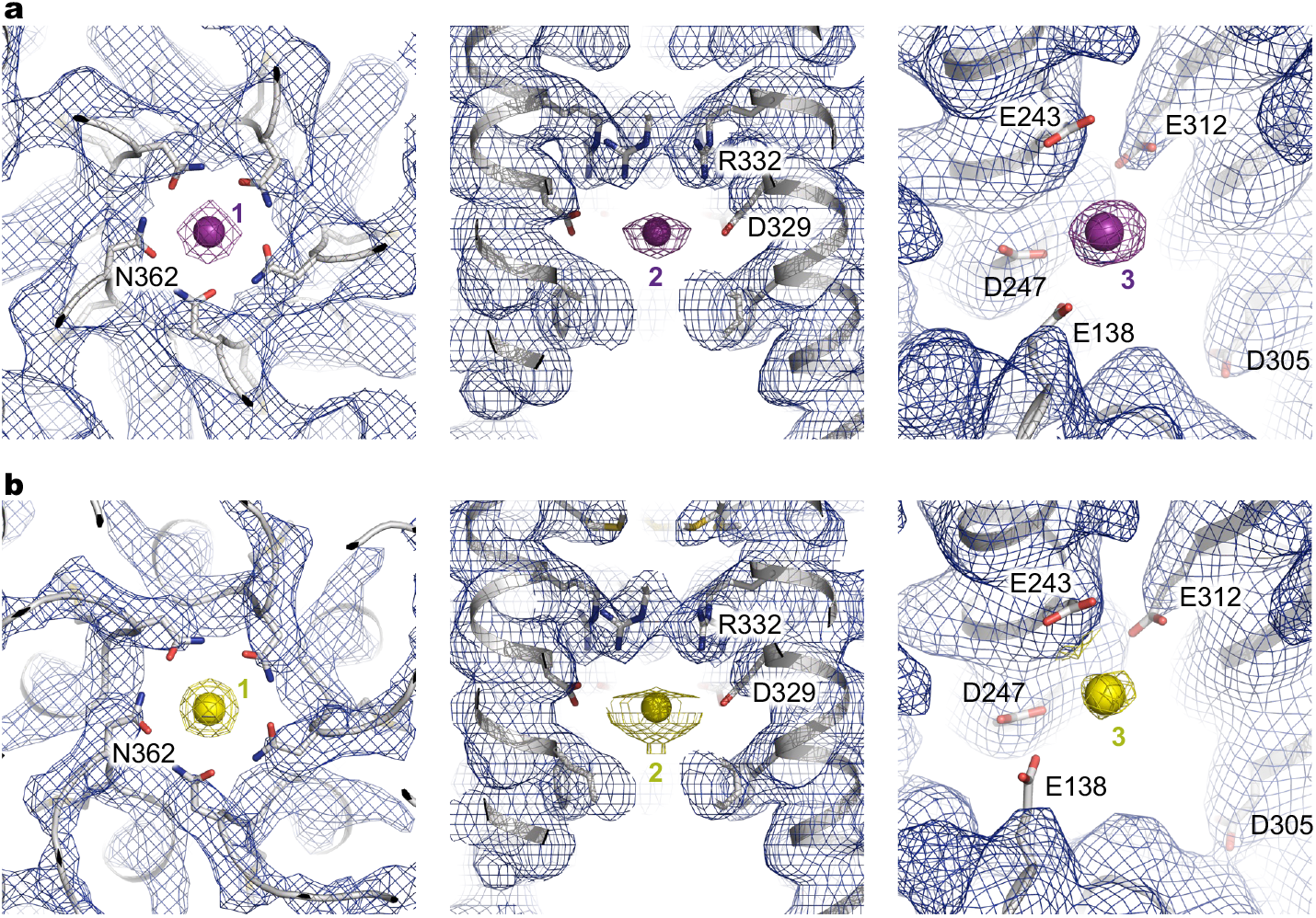
Ion densities. a-b, Cryo-EM densities in the three ion binding sites for Mg^2+^ (a) and Ca^2+^ (b). Mg^2+^ and Ca^2+^ are shown as magenta and yellow spheres, respectively.

**Extended Data Table 1.** Cryo-EM data collection, refinement and validation statistics.

|  | MRS2<sub>Mg</sub><br>(EMD-41587)<br>(PDB 8TS1) | MRS2<sub>EDTA</sub><br>(EMD-41588)<br>(PDB 8TS2) | MRS2<sub>Ca</sub><br>(EMD-41589)<br>(PDB 8TS3) |
| --- | --- | --- | --- |
| **Data collection and processing** |  |  |  |
| Magnification | 120k | 120k | 64k |
| Voltage (kV) | 200 | 200 | 300 |
| Electron exposure (e<sup>-</sup>/Å<sup>2</sup>) | 41 | 49 | 52 |
| Defocus range (μm) | -1.0 to -2.4 | -1.0 to -2.4 | -0.8 to -2.5 |
| Pixel size (Å) | 1.2 | 1.2 | 1.1 |
| Symmetry imposed | C5 | C5 | C5 |
| Initial particle images (no.) | 937,963 | 1,249,448 | 582,296 |
| Final particle images (no.) | 719,778 | 406,358 | 125,279 |
| Map resolution (Å) | 2.92 | 2.95 | 3.11 |
| FSC threshold | 0.143 | 0.143 | 0.143 |
| Map resolution range (Å) | 2.5-4.5 | 2.5-4.5 | 2.5-4.5 |
| **Refinement** |  |  |  |
| Initial model used (PDB code) | AlphaFold | This study | This study |
| Model resolution (Å) | 3.1 | 3.1 | 3.5 |
| FSC threshold | 0.5 | 0.5 | 0.5 |
| Map sharpening *B* factor (Å<sup>2</sup>) | -100 | -100 | -60 |
| Model composition |  |  |  |
| Nonhydrogen atoms | 12,202 | 12,240 | 12,027 |
| Protein residues | 1,505 | 1,505 | 1,505 |
| Ligands | MG: 7 | 0 | CA: 7 |
| *B* factors (Å<sup>2</sup>) |  |  |  |
| Protein | 78.12 | 114.31 | 143.27 |
| Ligand | 37.14 | N/A | 30.00 |
| R.m.s. deviations |  |  |  |
| Bond lengths (Å) | 0.003 | 0.004 | 0.002 |
| Bond angles (°) | 0.463 | 0.453 | 0.451 |
| Validation |  |  |  |
| MolProbity score | 1.12 | 1.09 | 1.17 |
| Clash score | 3.25 | 3.04 | 3.86 |
| Poor rotamers (%) | 0.37 | 0.73 | 0 |
| Ramachandran plot |  |  |  |
| Favored (%) | 98.64 | 98.98 | 98.64 |
| Allowed (%) | 1.36 | 1.02 | 1.36 |
| Disallowed (%) | 0 | 0 | 0 |

